# High Throughput Screening Of FDA Approved Drugs As Potential Agonist Of Mu-Opioid Receptor

**DOI:** 10.64898/2026.11.24.769065

**Authors:** Michael Olawale Olaoyo, Ruth Temitope Abioye, Tomilola Deborah Adeyinka

**Author notes:** is the corresponding author.

## Abstract

The mu-opioid receptor (MOR) remains an important analgesic target, but the interpretation of ligand-directed signalling preferences requires discrimination between receptor-dependent responses and effects on downstream assay readouts. We screened a library of 1,875 drug compounds for inhibition of forskolin-stimulated cyclic AMP (cAMP) accumulation in Chinese hamster ovary cells expressing human MOR and a GloSensor reporter. Primary screening of compound mixtures identified 17 active wells containing 204 compounds. Deconvolution yielded nine initial hits, six of which produced interpretable concentration–response relationships: niclosamide, sanguinarine, hexachlorophene, ataluren, altrenogest and nitazoxanide. Five of the six showed maximal cAMP inhibition of 77–80% in the initial concentration–response experiment, compared with 81 ± 6% for DAMGO; sanguinarine had a lower maximum (24 ± 6%). All six were less potent than DAMGO. In antagonist experiments, 300 nM naloxone shifted DAMGO potency from 0.3 to 224 nM. Of the tested compounds, only altrenogest showed a statistically significant naloxone-associated shift (367 to 905 nM; P = 0.0117). Altrenogest produced no detectable β-arrestin2 recruitment under the PathHunter assay conditions, whereas DAMGO produced a concentration-dependent response. Thus, most cAMP hits did not show evidence of naloxone-sensitive MOR activation, and altrenogest showed a limited but measurable antagonist-sensitive component. These results highlight the importance of orthogonal receptor-dependence testing before assigning MOR agonism or signalling bias to repurposing hits.

## Introduction

Pain is a multidimensional sensory and emotional experience and persistent pain represents a substantial clinical burden (Raja et al., 2020; Treede et al., 2019). Opioid analgesics are effective in selected settings, but adverse effects, tolerance and dependence constrain their use, particularly during prolonged treatment (Benyamin et al., 2008; Volkow and McLellan, 2016). Identifying analgesic mechanisms that preserve therapeutic activity while reducing unwanted effects remains an important goal of pharmacological research.

MOR is a G protein-coupled receptor that predominantly couples to inhibitory Gαi/o proteins. Receptor activation can suppress adenylyl cyclase activity and intracellular cAMP accumulation, while Gβγ-mediated modulation of ion channels can reduce neuronal excitability and transmitter release (Al-Hasani and Bruchas, 2011). Agonist-induced receptor phosphorylation also promotes β-arrestin recruitment, which can contribute to receptor trafficking, desensitisation and other signalling events. These multiple outputs motivated the hypothesis that ligands favouring G-protein-mediated signalling over β-arrestin recruitment might have improved therapeutic profiles.

However, the relationship between apparent pathway bias and clinical safety is not straightforward. Differences in intrinsic efficacy, receptor reserve, assay amplification and cellular context can generate apparent signalling preferences even when the underlying ligand mechanism is not genuinely pathway-selective (Gillis et al., 2020; Kliewer et al., 2019; Singleton et al., 2021). Moreover, a compound that lowers cAMP in MOR-expressing cells need not act at MOR: endogenous receptors, adenylyl cyclase, phosphodiesterases, cell viability or reporter chemistry may contribute to the observed signal. Consequently, receptor-dependence controls are essential before interpreting cAMP inhibition as MOR agonism.

Repurposing libraries offer an efficient way to explore pharmacologically characterised chemical space. Here, 1,875 compounds from an FDA-drug library were examined using a GloSensor cAMP assay, followed by deconvolution, concentration–response analysis, naloxone antagonism and a PathHunter β-arrestin2 recruitment assay. The study was designed to distinguish primary cAMP-active compounds from candidates with evidence of MOR-dependent activity and to assess whether the strongest receptor-sensitive hit displayed an apparent signalling preference under the assay conditions.

## Materials and methods

### Reagents and cellular model

PathHunter Chinese hamster ovary (CHO) cells and proprietary chemiluminescent assay reagents were obtained from DiscoverX, while luciferin potassium salt and pGloSensor-22F cDNA were purchased from Promega. Cell culture media, supplements, and assay buffer components were obtained from Gibco. The library of FDA-approved compounds was provided by the Drug Discovery Unit (DDU), School of Life Sciences, University of Dundee, UK. PathHunter CHO cells stably expressing human μ-opioid receptors (MOR) and β-arrestin2 components of the β-galactosidase enzyme fragment complementation system were used to assess intracellular cyclic adenosine monophosphate (cAMP) accumulation and β-arrestin2 recruitment. The same cell line was employed for both assays to maintain a consistent cellular background and receptor expression profile. Cells were maintained in Dulbecco’s Modified Eagle Medium (DMEM) supplemented with fetal bovine serum (FBS), penicillin (5,000 U/mL), and streptomycin (5,000 μg/mL). Geneticin (500 μg/mL) and hygromycin B (250 μg/mL) were included in the culture medium to maintain selection pressure for stable expression of the MOR and β-arrestin2 assay components. Cells were cultured at 37 °C in a humidified atmosphere containing 5% CO_2_ and routinely passaged upon reaching approximately 80% confluence.

### GloSensor transfection and cAMP accumulation assay

Cells were plated at 1 × 10^6^ per 35-mm dish in reduced-serum medium and transfected the following day with 2 μg pGloSensor-22F plasmid using Lipofectamine 2000 at a DNA-to-reagent ratio of 1:2.5. Transfected cells were incubated for at least 16 h; a parallel GFP-transfected control dish was assessed by fluorescence microscopy. Cells were subsequently seeded into half-area 96-well plates at 1.5 × 10^4^ cells per well and allowed to settle overnight. The culture medium was replaced with HBSS-based assay buffer containing 20 mM HEPES, 3 mM luciferin and 30 μM forskolin at pH 7.4. After 2 h at room temperature in darkness, baseline luminescence was measured using a GloMAX Navigator (1-s integration). Compounds or DAMGO, prepared at 5× working concentration, were added; following 30 min at room temperature, luminescence was measured again. The response was expressed relative to forskolin-stimulated baseline luminescence.

### Library screening and hit deconvolution

The compounds were supplied in twenty 96-well plates as 500 μM stock solutions in dimethyl sulfoxide (DMSO) and stored at −80 °C until use. For high-throughput screening, the twenty drug plates were organised into four screening sets comprising 480, 480, 480 and 435 compounds, respectively. Compounds were screened at a final concentration of 1 μM using a plate-consolidation strategy in which twelve compounds were pooled into individual assay wells. For plate preparation, intermediate V-bottom 96-well plates were prepared by dispensing 44 μL of assay working solution into designated wells, followed by the addition of 0.5 μL of each compound stock. Subsequently, 10 μL of the prepared compound mixtures was transferred into the corresponding cell assay wells for screening in duplicate. DAMGO (1 μM) served as the positive control, while the vehicle control consisted of Hank’s Balanced Salt Solution (HBSS) supplemented with 20 mM HEPES, 3 mM luciferin (Promega, UK) and 30 μM forskolin (HelloBio, UK), adjusted to pH 7.4, with an equivalent volume of DMSO to match the vehicle concentration in compound-treated wells. Compound mixtures producing responses comparable to DAMGO were selected for secondary screening, during which the twelve individual compounds within each active pool were tested separately to identify those responsible for the observed inhibition of forskolin-stimulated cAMP accumulation.

### Naloxone antagonism

To determine whether the inhibition of forskolin-stimulated cAMP accumulation by the identified compounds was mediated through μ-opioid receptor (MOR) activation, a naloxone antagonism assay was performed. Naloxone, a competitive opioid receptor antagonist, was used to assess the receptor dependence of the observed responses and distinguish MOR-mediated activity from potential off-target effects. Briefly, CHO cells were treated with assay working solution, followed by the addition of naloxone (300 nM), prepared in the same working solution. The test compounds were subsequently added, and the cells were incubated for 30 min at 37 °C. Following incubation, luminescence was measured using a GloMax Navigator luminometer to assess changes in intracellular cAMP accumulation. Concentration–response relationships were generated for each compound in the presence and absence of naloxone to evaluate changes in potency (IC_50_) and maximal inhibitory efficacy (Emax).

### β-arrestin2 recruitment

For the PathHunter enzyme-fragment complementation assay, cells were resuspended in Opti-MEM and seeded into half-area 96-well plates at 5 × 10^3^ cells per well. After overnight incubation at 37 °C, compounds prepared at 5× concentrations in Opti-MEM were added for 90 min at 37 °C. Chemiluminescent substrate was added, followed by 2 h at room temperature in darkness, and luminescence was measured on a GloMAX Navigator with 1-s integration. DAMGO served as the positive reference ligand. Altrenogest was selected for this follow-up because its cAMP response showed significant naloxone sensitivity.

### Data analysis and statistics

Luminescence values obtained from the β-arrestin2 recruitment assay were normalised and expressed as a percentage of the maximum relative luminescence units (RLU) recorded on each plate, while responses from the cAMP accumulation assay were expressed as a percentage of forskolin-stimulated luminescence measured before compound addition. For naloxone antagonism experiments, responses were normalised to the maximal agonist-induced response in the absence of naloxone and expressed as a percentage of control. Agonist potency and efficacy were compared with those of the reference MOR agonist, DAMGO, except in naloxone antagonism experiments, where each compound served as its own control. Concentration–response relationships were analysed using GraphPad Prism and fitted to a three-parameter logistic equation:

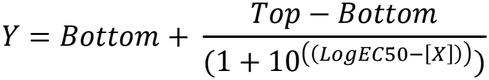

where Y represents the measured response, X is the logarithm of the agonist concentration, and Top and Bottom represent the maximum and minimum fitted responses, respectively. The equation was applied using EC_50_ for β-arrestin2 recruitment and IC_50_ for inhibition of cAMP accumulation, as appropriate. Individual concentration–response datasets were fitted independently to determine agonist potency (EC_50_ or IC_50_) and maximal efficacy (Emax), calculated as the difference between the fitted maximum and minimum responses. Potency estimates were transformed into pEC_50_ or pIC_50_ values, defined as the negative base-10 logarithm of the corresponding molar concentration. The minimum response was constrained to values ≥ 0 for both assays. Mean concentration–response curves were also fitted using the same logistic model for graphical presentation only, while statistical analyses were performed using parameters derived from individual fitted datasets. Differences in agonist efficacy (Emax) and potency (pEC_50_ or pIC_50_) relative to DAMGO were assessed using one-way analysis of variance (ANOVA), followed by Bonferroni’s multiple-comparison test. Data were assessed for normality and homogeneity of variance before parametric analysis, and non-parametric tests were applied when these assumptions were not satisfied, including after data transformation. Statistical significance was defined as p < 0.05. Summary data are presented as mean ± standard error of the mean (SEM), with potency estimates reported alongside 95% confidence intervals, as appropriate.

## Results

### Primary screening and compound deconvolution

In the primary screen, 1,875 compounds were evaluated as twelve-compound mixtures in four assay plates. Wells whose responses approached the DAMGO positive-control region were advanced, whereas mixtures resembling vehicle controls were not. Seventeen primary wells met the selection criterion, representing 204 candidate compounds. Each constituent was then tested individually. Nine compounds retained a positive cAMP-screening signal, but three were excluded because concentration–response relationships were inconsistent. Six compounds therefore proceeded to quantitative characterisation (Figures 1 and 2). Because the initial readout measures cAMP-associated luminescence, these hits are described as cAMP-active compounds rather than confirmed MOR agonists.

**Figure 1.**
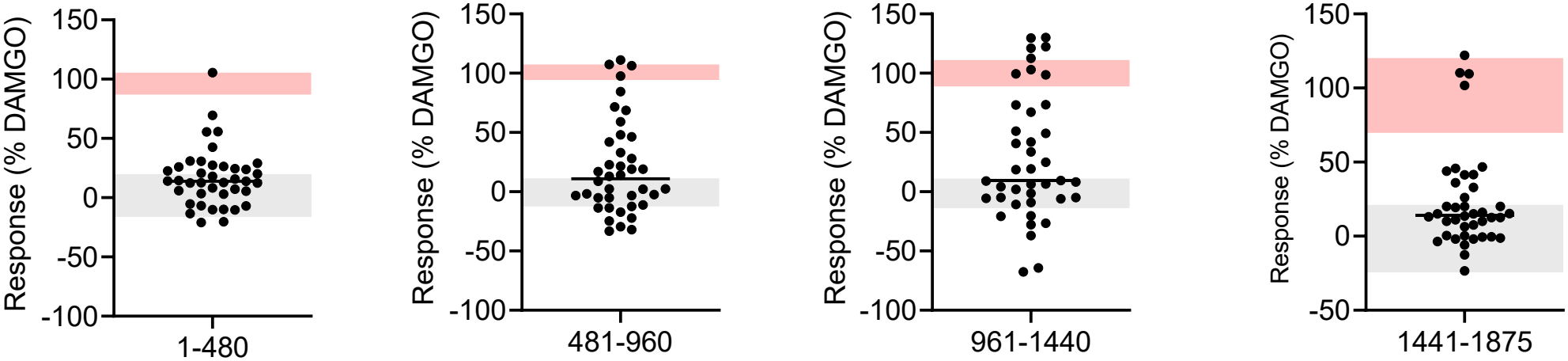
Primary screening of 1,875 compounds in four groups. Points represent compound-mixture responses relative to DAMGO; shaded regions indicate the DAMGO-like and vehicle-like selection bands.

**Figure 2.**
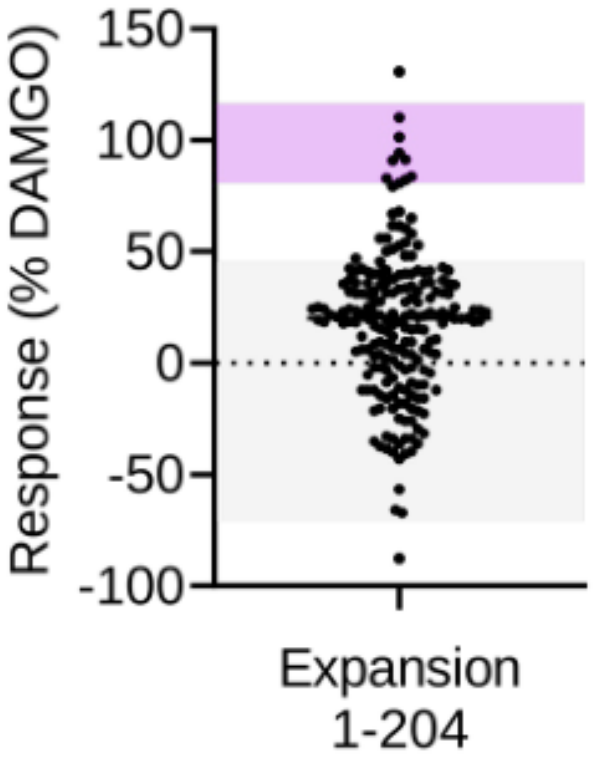
Secondary deconvolution of 204 individual compounds from the seventeen selected mixture wells.

**Figure 3.**
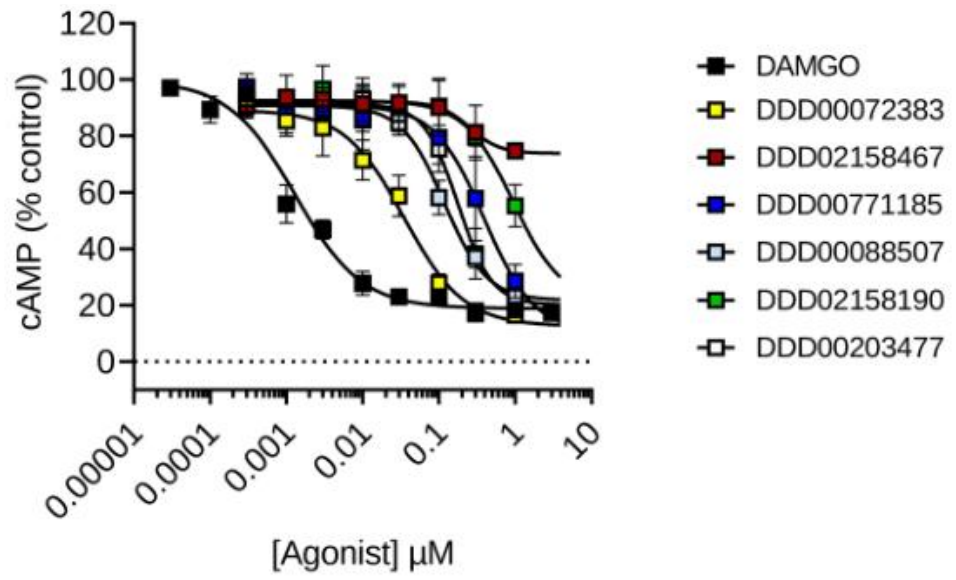
Concentration–response curves for DAMGO and the six selected compounds in the GloSensor cAMP assay.

**Figure 4.**
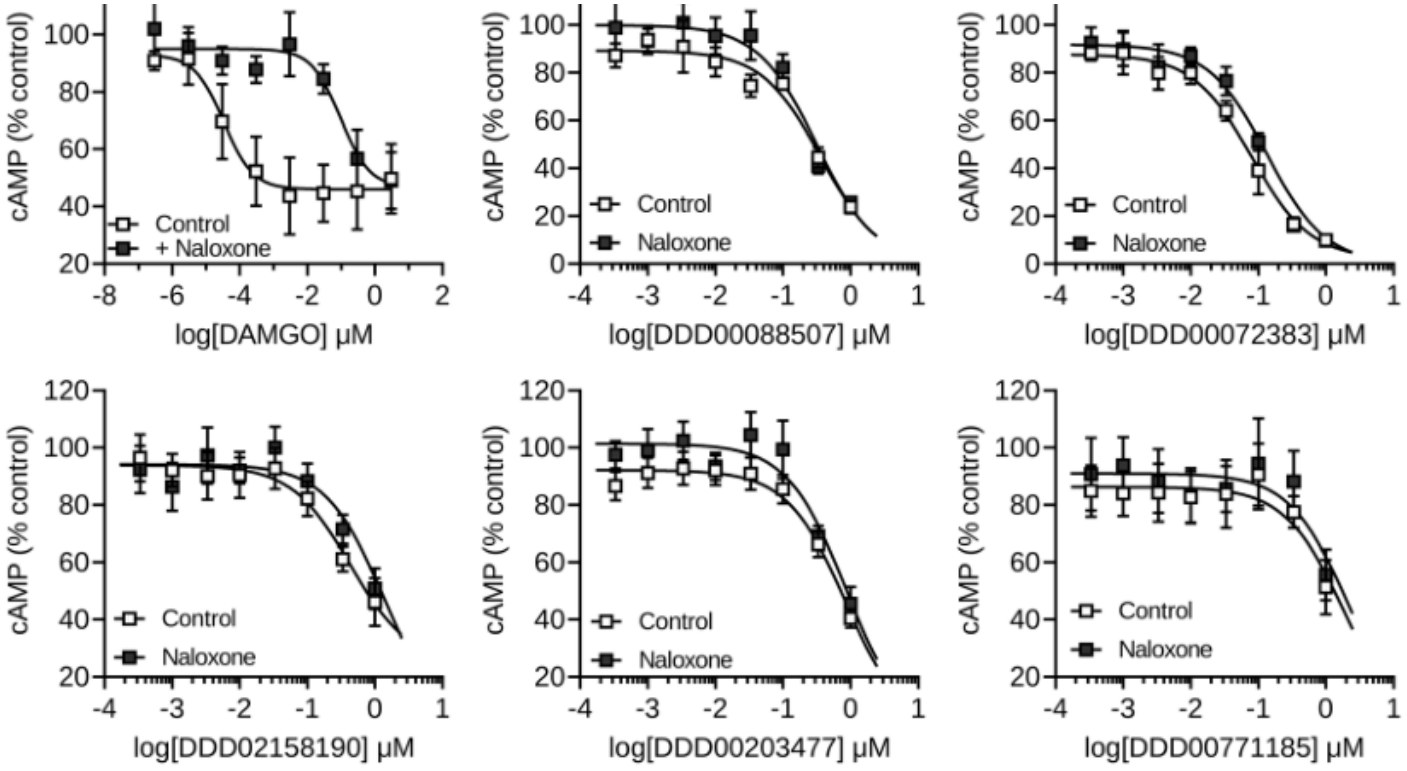
Concentration–response comparisons for DAMGO and selected compounds with or without 300 nM naloxone. Original figure reports mean ± SEM from five independent experiments.

### Six hits show concentration-dependent cAMP inhibition

Niclosamide (DDD00072383), sanguinarine (DDD02158467), hexachlorophene (DDD00771185), ataluren (DDD00088507), altrenogest (DDD02158190) and nitazoxanide (DDD00203477) yielded fitted concentration–response curves. DAMGO produced an Emax of 81 ± 6% and an IC_50_ of 1 nM (95% CI, 1–2 nM). Niclosamide, hexachlorophene, ataluren, altrenogest and nitazoxanide showed similar maximal cAMP inhibition (77–80%), whereas sanguinarine showed a substantially smaller maximum (24 ± 6%; P < 0.001). All six compounds were less potent than DAMGO. Among the hits, niclosamide had the lowest reported IC_50_ (36 nM, 95% CI 26–47), whereas altrenogest had the highest (1,415 nM, 95% CI 1,052–2,252). The reported overall group comparisons were significant for both efficacy (F_6,20_ = 24.2, P < 0.0001) and potency (F_6,20_ = 35.5, P < 0.0001). These findings established activity in the cAMP assay but did not independently establish MOR engagement.

### Naloxone identifies a receptor-antagonist-sensitive component for altrenogest

Naloxone (300 nM) caused a pronounced rightward displacement of the DAMGO concentration– response curve: IC_50_ increased from 0.3 nM (95% CI 0.02–0.57) to 224 nM (95% CI 50–468), while Emax remained similar (48 ± 13% versus 50 ± 12%). The potency difference was significant (t = 8.130, df = 8, P < 0.0001), but the efficacy difference was not (P = 0.9069), consistent with competitive antagonism of the reference agonist.

In contrast, naloxone did not significantly alter the cAMP potency or efficacy of niclosamide, hexachlorophene, ataluren or nitazoxanide. Niclosamide IC_50_ changed from 115 to 131 nM (P = 0.2588), hexachlorophene from 2,584 to 2,170 nM (P = 0.9532), ataluren from 365 to 393 nM (P = 0.8242), and nitazoxanide from 1,093 to 1,121 nM (P = 0.958). These data do not support substantial naloxone-sensitive MOR-mediated activity for these four compounds under the tested conditions.

Altrenogest differed from the other hits: its IC_50_ increased from 367 nM (95% CI 203–541) to 905 nM (95% CI 326–1,525) with naloxone (t = 3.381, P = 0.0117). Emax was not significantly changed (65 ± 13% versus 76 ± 11%; P = 0.1710). The approximately 2.5-fold potency shift was considerably smaller than that observed with DAMGO, suggesting a naloxone-sensitive component of the altrenogest response but not establishing its precise binding site or mechanism.

### Altrenogest does not detectably recruit β-arrestin2 in the assay

Following the naloxone antagonism experiments, altrenogest (DDD02158190) was selected for further investigation because it was the only test compound that exhibited a significant naloxone-sensitive reduction in potency, suggesting involvement of the μ-opioid receptor (MOR). To determine whether altrenogest also promoted β-arrestin2 recruitment, its activity was evaluated using the PathHunter assay in CHO cells stably overexpressing human MOR. Concentration– response relationships were generated for altrenogest and the reference MOR agonist DAMGO, as shown in Figure 5. DAMGO induced a concentration-dependent increase in β-arrestin2 recruitment, with a maximal efficacy (Emax) of 96 ± 4% and an EC_50_ of 423 nM (95% CI: 170– 807 nM), as presented in Table 3. In contrast, altrenogest produced no detectable β-arrestin2 recruitment across the concentrations tested, preventing the determination of its maximal efficacy and potency (ND). These findings demonstrate that, under the experimental conditions employed, altrenogest exhibited naloxone-sensitive inhibition of cAMP accumulation without measurable β-arrestin2 recruitment, suggesting a potential preference for G-protein-mediated signalling over β-arrestin2 recruitment at MOR.

**Table 1.** Efficacy and potency estimates for inhibition of forskolin-stimulated cAMP accumulation.

| **Compounds** | **EMAX (%)** | **IC50 nM (95% CI)** |
|---|---|---|
| DAMGO | 81 ± 6% | 1 [1-2] |
| DDD00072383 (NICLOSAMIDE) | 80 ± 2% | 36 [26-47]*** |
| DDD02158467 (SANGUINARINE) | 24 ± 6%*** | 290 [11-587]*** |
| DDD00771185(HEXACHLOROPHENE) | 77 ± 4% | 346 [286-411]*** |
| DDD00088507 (ATALUREN) | 77 ± 3% | 146 [109-171]*** |
| DDD02158190 (ALTRENOGEST) | 77 ± 3% | 1415 [1052-2252]*** |
| DDD00203477 (NITAZOXANIDE) | 79 ± 1% | 223 [148-392]*** |

**Table 2.** Naloxone antagonism: original efficacy and potency estimates.

| **Compound** | **Condition** | **EMAX (Mean ± SEM)** | **Potency (nM) 95% CI** |
|---|---|---|---|
| DAMGO | Control | 48 ± 13 | 0.3 [0.02 – 0.57] |
|  | Naloxone (300nM) | 50 ± 12** | 224 [50 – 468] |
| DDD00072383 (NICLOSAMIDE) | Control | 87 ± 5 | 115 [45 – 196] |
|  | Naloxone | 93 ± 6 | 131 [98 – 176] |
| DDD00771185 (HEXACHLOROPHENE) | Control | 86 ± 9 | 2584 [1100 – 4476] |
|  | Naloxone | 91 ± 11 | 2170 [1771 – 2512] |
| DDD02158190 (ALTRENOGEST) | Control | 65 ± 13 | 367 [203 – 541] |
|  | Naloxone | 76 ± 11** | 905 [326 – 1525] |
| DDD000888507 (ATALUREN) | Control | 86 ± 5 | 365 [220 – 535] |
|  | Naloxone | 97 ± 8 | 393 [243 – 567] |
| DDD00203477 (NITAZOXANIDE) | Control | 90 ± 5 | 1093 [452 – 1831] |
|  | Naloxone | 95 ± 8 | 1121 [443 – 1856] |

**Table 3.** β-arrestin2 recruitment efficacy and potency (ND, not determined).

| **Compounds** | **EMAX (%)** | **EC<sub>50</sub> nM (95% CI)** |
|---|---|---|
| DAMGO | 96 ± 4% | 423 [170-807] |
| DDD02158190 | ND | ND |
| (ALTRENOGEST) |  |  |

**Figure 5.**
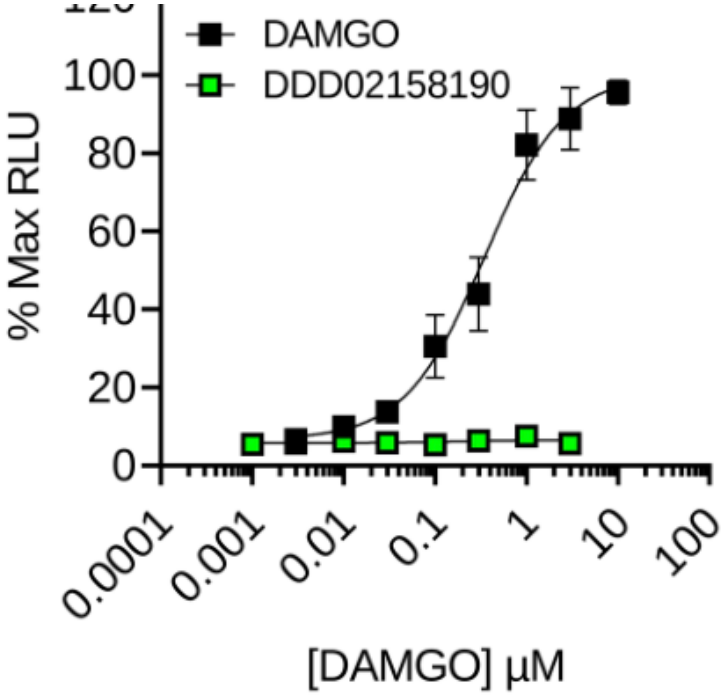
PathHunter β-arrestin2 recruitment responses to DAMGO and altrenogest. DAMGO produced concentration-dependent recruitment; altrenogest showed no quantifiable response.

### Cross-pathway comparison is descriptive rather than a formal bias estimate

The relative signalling profiles of DAMGO and altrenogest were evaluated by comparing their maximal inhibitory effects on forskolin-stimulated cAMP accumulation with their β-arrestin2 recruitment responses. As altrenogest produced no quantifiable β-arrestin2 concentration–response relationship, its apparent efficacy was estimated from the difference between responses at the highest and lowest concentrations tested. These values were plotted against the corresponding maximal cAMP inhibitory responses to assess apparent functional selectivity (Figure 6). DAMGO exhibited substantial efficacy in both signalling assays, whereas altrenogest demonstrated pronounced cAMP inhibition with minimal β-arrestin2 recruitment. This divergence in signalling profiles suggests an apparent preference for cAMP-associated signalling over β-arrestin2 recruitment by altrenogest under the experimental conditions tested.

**Figure 6.**
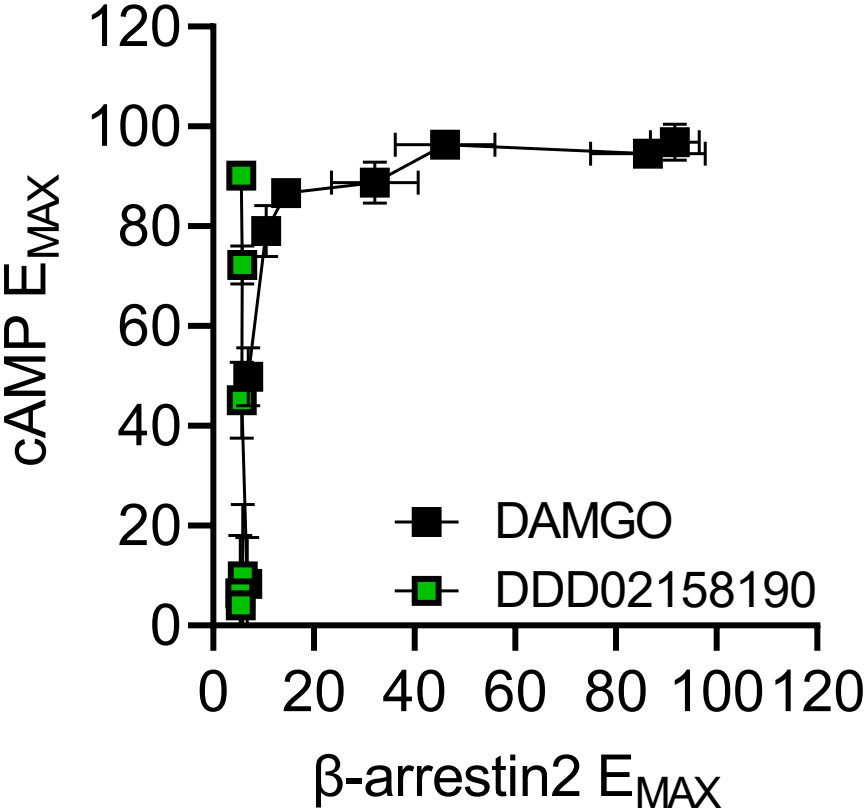
Descriptive comparison of maximal cAMP inhibition and β-arrestin2 recruitment for DAMGO and altrenogest. This plot does not constitute a quantitative bias-factor calculation.

## Discussion

The central finding of this screen is that cAMP-active compounds identified in MOR-expressing cells are not necessarily MOR agonists. Screening 1,875 drug-library compounds yielded six hits with interpretable concentration–response relationships, yet only altrenogest displayed a statistically significant naloxone-associated reduction in apparent potency. This separation between primary functional activity and antagonist sensitivity is particularly important for repurposing campaigns, where compounds may interact with numerous cellular pathways or assay components.

The first stage of screening was intentionally sensitive to compounds that reduced forskolin-stimulated cAMP-associated luminescence. That strategy allowed the rapid triage of a large chemical collection, but the measured endpoint is downstream of MOR and does not uniquely report receptor occupancy. The five hits with large cAMP responses in the initial curves illustrate the potential for downstream assay activity to resemble DAMGO efficacy. Their generally lower potency than DAMGO and the absence of substantial naloxone-induced shifts in four of the compounds caution against treating maximal cAMP inhibition as evidence of opioid agonism.

The DAMGO antagonist experiment provided an internal pharmacological benchmark. Naloxone caused a large rightward potency shift with little change in maximum response, as expected for competitive antagonism under the assay conditions. By comparison, altrenogest showed a smaller, approximately 2.5-fold rightward shift. This observation supports the possibility that some component of its cAMP response is opioid-receptor-sensitive. Nevertheless, a modest antagonist shift alone cannot demonstrate direct orthosteric binding, distinguish full from partial receptor involvement or rule out compound-dependent interference with the reporter system. The absence of orthogonal target-engagement measurements remains a limitation.

Altrenogest did not induce detectable β-arrestin2 recruitment in the PathHunter system. Considered alongside the cAMP data, this pattern suggests an apparent difference between the two readouts, but it should not be equated with a validated ligand-bias phenotype. Receptor expression, pathway amplification, signal window, compound solubility and assay sensitivity can influence whether a low-efficacy response is measurable. Recent work on MOR agonists underscores that apparent bias may be confounded by intrinsic efficacy and receptor reserve (Gillis et al., 2020; Kliewer et al., 2019; Singleton et al., 2021).

For the naloxone-insensitive compounds, several alternative mechanisms remain plausible. They may affect endogenous signalling pathways in CHO cells, alter adenylyl cyclase or phosphodiesterase activity, perturb cellular metabolism or directly interfere with luciferase-dependent signal production. The current experiments cannot discriminate among these explanations. In particular, the absence of naloxone sensitivity does not identify a specific alternative GPCR, and it does not completely exclude noncanonical interactions with MOR. Additional counterscreens in parental cells lacking MOR, orthogonal cAMP assays and cell-viability or reporter-interference controls would be needed to determine which compounds act on cAMP biology rather than the optical readout.

The next steps would be to repeat altrenogest concentration–response measurements in independently prepared cultures, assess antagonist concentration dependence, confirm activity using an orthogonal cAMP measurement, and compare responses in MOR-positive and MOR-negative cells. Pertussis-toxin sensitivity could help assess Gi/o involvement, while additional β-arrestin recruitment methods and direct target-engagement assays would test the robustness of the signalling phenotype. These studies would clarify whether altrenogest engages MOR directly and whether any pathway preference persists after controlling for assay amplification and receptor reserve.

In conclusion, this study demonstrates the value of combining high-throughput functional screening with pharmacological antagonist controls. Most apparent cAMP hits were not naloxone-sensitive, while altrenogest showed a limited but significant antagonist-sensitive cAMP response without measurable β-arrestin2 recruitment. The results identify a candidate for mechanistic follow-up rather than a validated biased MOR agonist or a proven safer analgesic.

## Declarations

## Acknowledgements

The authors gratefully acknowledge Professor Tim Hales and the School of Life Sciences at the University of Dundee for their support and contributions to this research.

## Author Contributions

Michael Olawale Olaoyo: Conceptualisation, methodology, investigation, formal analysis, data curation, visualisation, and writing – original draft.

Ruth Temitope Abioye: Literature review, interpretation of findings, and writing – review and editing.

Tomilola Deborah Adeyinka: Interpretation of findings, critical review, and writing – review and editing.

All authors read and approved the final manuscript and agreed to its submission.

